# Wavelet Screening: a novel approach to analysing GWAS data

**DOI:** 10.1101/2020.03.24.006163

**Authors:** William Denault, Håkon K. Gjessing, Julius Juodakis, Bo Jacobsson, Astanand Jugessur

**Affiliations:** Department of Genetics and Bioinformatics, Norwegian Institute of Public Health, Oslo, Norway; Centre for Fertility and Health, Norwegian Institute of Public Health, Oslo, Norway; Department of Global Public Health and Primary Care, University of Bergen, Bergen, Norway; Department of Obstetrics and Gynecology, Institute of Clinical Sciences, Sahlgrenska Academy, University of Gothenburg, Gothenburg, Sweden

**Keywords:** SNP, GWAS, Multiple testing, Polygenic association, Wavelet, Wavelet regression

## Abstract

We present here an alternative method for genome-wide association study (GWAS) that is more powerful than traditional GWAS methods for locus detection. Single-variant GWAS methods incur a substantial multiple-testing burden because of the vast number of single nucleotide polymorphisms (SNPs) being tested simultaneously. Furthermore, these methods do not consider the functional genetic effect on the outcome because they ignore more complex joint effects of nearby SNPs within a region. By contrast, our method reduces the number of tests to be performed by screening the entire genome for associations using a sliding-window approach based on wavelets. In this context, a sequence of SNPs represents a genetic signal, and for each screened region, we transform the genetic signal into the wavelet space. The null and alternative hypotheses are modelled using the posterior distribution of the wavelet coefficients. We enhance our decision procedure by using additional information from the regression coefficients and by taking advantage of the pyramidal structure of wavelets. When faced with more complex signals than single-SNP associations, we show through simulations that Wavelet Screening provides a substantial gain in power compared to both the traditional GWAS modelling as well as another popular regional-based association test called ‘SNP-set (Sequence) Kernel Association Test’ (SKAT). To demonstrate feasibility, we re-analysed data from the large Norwegian HARVEST cohort.

## 1. Introduction

The objective of a genetic association study is to identify genes and loci that are associated with a given phenotype of interest. Although the human genome is very similar across individuals, it is interspersed with single base-pair differences called single-nucleotide polymorphisms (SNPs) that collectively contribute to the observed differences across individuals. One of the most common approaches to screening for genetic associations with a trait or disorder is to conduct a genome-wide association study (GWAS) where the significance of the effect of each SNP is assessed in a sequential fashion. Despite its many successes, however, this approach is limited due to two important issues: (i) it incurs a substantial multiple-testing burden because of the large number of tests carried out simultaneously, and (ii) it ignores the functional nature of the genetic effect by failing to exploit the dense genotyping of the genome. Because larger regions of the genome might contribute to the phenotype, only considering the effect of one SNP at a time would not efficiently model how a larger change in the genome might contribute to the phenotype.

The issue of multiple testing can be resolved using a regularisation method such as Fused Lasso (Robert Tibshirani and Knight, 2005). The main idea behind Fused Lasso is to perform a penalised regression that takes into account how variables (here, SNPs) that are physically close to each other might produce similar effects. Using Fused Lasso one may define a region of association between the SNPs and the phenotype. However, Fused Lasso only performs local testing, whereas it is more desirable to test for associations in larger chunks of the genome.

Despite an increased interest in penalised regressions within the broader statistical community, they rarely appear in the top-tiered genetic publications. Penalised regression has recently been incorporated in one of the leading software for GWAS known as PLINK (Purcell *and others*, 2007), but the lack of a comparable software for meta-analysis is a major limitation of this approach. This is primarily because a comprehensive genome-wide association meta-analysis (GWAMA) typically relies on analysing summary statistics from multiple cohorts. Even though meta-analyses are now feasible in the Lasso regression setting (Lockhart *and others*, 2014), they are not currently available for variants of Lasso regression or for other regularisation penalties.

To address these shortcomings, we developed “Wavelet Screening” as a new approach to GWAS by harnessing insights from functional modeling (Morris and Carroll, 2006; Shim and Stephens, 2015). Specifically, we adapt the approach described by Shim and Stephens (2015) in which the authors test for association with a functional phenotype (the response signal) by first transforming the signal using fast discrete wavelet transform (Mallat, 2008) and then testing for single-SNP associations. In essence, our main idea is to reverse this approach, by modeling the SNP signals as wavelets over a large genomic region and regressing the wavelet coefficients on the phenotype.

The use of reverse regression to search for associations is now more widespread in the genetic literature (see (Aschard *and others*, 2017) for an example). Our approach provides a broader test by enabling an estimation of the fraction of wavelet coefficients (blocks) associated at each level of depth, and by treating sizable chunks of the genome (*≈* 1 million base pairs) as the functional phenotype. We then perform a dimensional reduction using wavelet transform and test for associations between the wavelet coefficients and the phenotype. This broader approach to testing combined with multiple levels of information may provide additional insights into the mechanisms underlying a detected genetic association.

By reversing the regression and targeting a given region for association testing, we use regional association instead of single-SNP association to reduce the number of tests to be performed. Notably, by using overlapping windows of 1 Mb in length, one can reduce the number of tests to be performed from eight million (for common SNPs) to approximately 5000. We propose screening regions of 1 Mb in size in order to cover most of the linkage disequilibrium (LD) blocks in humans regardless of ethnicity (Wall and Pritchard, 2003). As our method provides region-wide p-values, it is straightforward to perform large-scale meta-analyses by simply combining p-values from different cohorts using Fisher’s method (Fisher, 1958).

The remainder of our paper is structured as follows. We first describe the statistical setting for the analyses and the wavelet methodology used to generate the wavelet coefficients. Next, we describe our test statistics between the wavelet spectrum and the phenotype Φ. The phenotype Φ in this paper represents a univariate vector of either a continuous, countable, or binary trait. After a comprehensive evaluation of the method by a series of simulations, we apply it to data from the Norwegian HARVEST study, which is a sub-project nested within the larger Norwegian Mother and Child Cohort Study (MoBa) (Magnus *and others*, 2016). Our prime phenotype of interest is gestational age at birth.

## 2. Description of Wavelet Screening

### 2.1 Haar wavelet transform

Our method transforms the raw genotypic data in a similar way to the widely used “Gene- or Region-Based Aggregation Tests of Multiple Variants” (Seunggeung Lee and Lin (2014)). Like the Burden test, the effects of the genetic variants in a given region are summed up to construct a genetic score for the regression. The first step in our analysis is to use fast discrete wavelet transform to transform the multi-SNP data. Next, we introduce the Haar wavelet transform and show how the wavelet coefficients are computed. Readers unfamiliar with wavelets are referred to a comprehensive introduction to wavelets by Nason (2008). In the rest of this article, ‘wavelet’ specifically refers to the Haar wavelet.

We code a SNP 0 if an individual is homozygous for the reference allele, 1 if heterozygous, and 2 if homozygous for the alternate allele, which is consistent with an additive genetic model and the standard way of coding alleles (Purcell *and others*, 2007).

Let *G*_0,*k*_(*bp*) denote the “true” genetic signal of individual *k* at physical position *bp* (base pair), and let *G*_*k*_(*bp*) be the observed, imputed version of *G*_0,*k*_(*bp*). We assume that

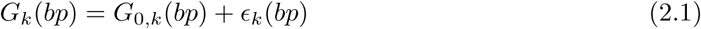

where *ϵ*_*k*_(*bp*) are independently and identically distributed over individuals, with Var(*ϵ*_k_(bp)) = *σ*^2^(bp). The variance *σ*^2^(*bp*) at position *bp* can be interpreted as a function of the imputation quality *IQ*(*bp*), which has a value in [0, 1]. 1 represents a perfectly imputed SNP or genotyped SNP; thus, *σ*^2^(*bp*) ∝ 1 − *IQ*(*bp*). As the data used here had already been quality controlled, only SNPs with an *IQ* ∈ [0.8, 1] are retained for further analyses. We assume that the imputation qualities are independent and heteroscedastic over *bp*. As the value of a SNP is in {0, 1, 2} and then in [0, 2] following the dosage convention after imputation (Purcell *and others*, 2007), the distribution of *ϵ*_*k*_(*bp*) is not straightforward. However, as our model is calibrated by simulations, this error distribution does not have to be specified.

We define a genetic region *GR*_*lb,ub*_ on a given chromosome as the set of physical positions *bp* in the interval *lb < bp < up*. For the rest of the paper, we assume the analyses are performed within a fixed genetic region *GR*_*lb,ub*_ on a given chromosome. We observe the value of *G*_*k*_(*bp*) at pre-determined and increasing positions *bp*_1_, …, *bp*_*n*_ within the interval (*lb, ub*), with some error due to the genome-wide imputation process (Li *and others*, 2009). For now, we assume having *n* = 2^*J*^ equally spaced observations within *GR*_*lb,ub*_ and denote the observed value of *G*_*k*_(*bp*_*i*_) by *g*_*k*_(*bp*_*i*_), i.e., the data value measured on individual *k* at position *bp*_*i*_, *i* = 1, *… n*, where the 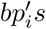 are equally spaced. We define wavelet *d* and *c* coefficients as sequences of length 2^*J*^. These coefficients are computed by Mallat’s pyramid algorithm (Mallat, 2008). For the coefficients at the highest scale (i.e., scale *J* − 1), for *i* ∈ {1, *…*, 2^*J*−1^},

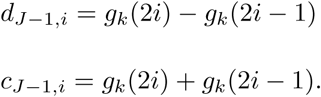

These coefficients correspond to local differences (or sums) of the measured values. For lower scales, the coefficients are computed as follows:

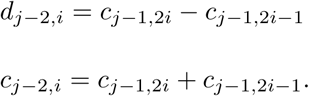

Finally, the coefficients at the lowest scale (i.e., scale 0) are computed as

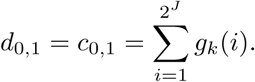

These procedures are often written as matrices *W*_*d*_ and *W*_*c*_ (d and c procedures, respectively), where the rows of *W*_*d*_ and *W*_*c*_ are normalised. We have *d* = *Wg*_*k*_ and *c* = *W*′*g*_*k*_. In addition, because the matrix *W*_*d*_ is orthogonal, we have:

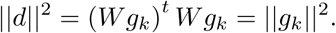

Using the 2^*J*^ wavelet coefficients for individual *k*, all values *g*_*k*_(*bp*) in the genetic region *GR*_*lb,ub*_ can be completely recovered. However, this wavelet transformation assumes the data to be evenly spaced, as well as *n* = 2^*J*^ measurements, which may not be realistic in practice. To avoid this assumption, we use the method of Kovac and Silverman (2000) which is briefly explained in Subsection 2.2.

#### Wavelet representation of the genom

Essentially, the coefficients obtained by performing the wavelet transform on a region of the genome can be viewed as local “scores” of the genotype, with the following interpretations:

- At scale 0, the wavelet coefficients *d* and *c* can be interpreted in the same way: they summarise the amount of discrepancy between an individual’s genotypes and the reference genotypes coded as 0…0. This is essentially the test comparison performed in standard gene or regional tests.
- The wavelet *d* coefficient at scale *s >* 0 and location *l* for an individual represents the difference in the number of minor alleles between the left part of the region (defined by *s, l*) and the right part.
- The wavelet *c* coefficient at scale *s >* 0 and location *l* for an individual represent the amount of discrepancy between an individual’s genotypes and the reference genotypes coded as 0…0 for the region defined by *s, l*.

The main rationale behind this modeling is that, if a genetic locus has an effect on the phenotype, the association is likely to be spread across genomic regions of a given size (scale) at different positions (locations). By using the wavelet transform to perform a position/size (time/frequency) decomposition and then regressing the wavelet coefficients on the phenotype, we are able to visualise *where* (location) and *how* (scale) the genetic signal influences the phenotype.

In the rest of this article, we use ‘wavelet coefficients’ to refer to *c* coefficients specifically. *c* coefficients are easier to interpret than *d* coefficients. For instance, in case of completely observed genotype, *c* coefficients correspond to the sum of minor alleles (similar to the Burden test (Ionita-Laza *and others*, 2013)).

### 2.2 Preprocessing of data

#### Non-decimated wavelet transfor

We use the method of Kovac and Silverman (2000) for non-decimated and unevenly spaced data. This method takes an irregular grid of data, for instance the sampling of different genetic regions, and interpolates the missing data into a pre-specified regular grid of length 2^*J*^. For a given genetic region *GR*_*lb,ub*_, with measurements at *n* positions *bp*_1_…*bp*_*n*_, we map this region into a (0, 1) interval using the affine transformation *x* 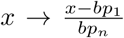. We then define a new grid of points of length 2^*J*^ on (0, 1) as: *t*_0_, …, *t*_*N*−1_, where *N* = 2^*J*^, *J* ∈ ℕ, 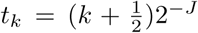 and *J* = *min*{*j* ∈ ℤ, 2^*j*^ ⩾ *n*}. We interpolate the mapped signal into this grid and run wavelet transform to obtain the wavelet coefficients. In practice (see Section 4), we recommend selecting genetic regions that have a relatively high density of imputed SNPs.

#### Coefficient-dependent thresholding and quantile transfor

For each individual wavelet decomposition, we use the VisuShrink approach (Kovac and Silverman, 2000) to shrink the interpolated wavelet coefficients and reduce the dependence between the wavelet coefficients within scales. This allows an estimation of the variance of each wavelet coefficient before determining a specific threshold for each wavelet coefficient. We can account for the individual heteroscedasticity of the noise by determining specific coefficient-dependent thresholds using the wavelet coefficient variance. Next, we quantile-transform each wavelet coefficient distribution within the population to make sure that each distribution follows a *N* (0, 1) distribution. Because we use the quantile-transformed wavelet coefficient as the endogenous variable (see Section 2.3), the above transformation ensures that, under the null hypothesis, the residuals are normally distributed. This also controls for spurious associations resulting from any deviation from the Normal distribution assumption of linear model.

### 2.3 Modelling

In essence our approach to modelling aims to detect regions that contain sub-regions associated with a trait of interest. In the second step, we localize these sub-regions to ease the interpretation of the output. We first need to assess whether certain scales are associated with the phenotype at different locations to estimate the effect of a genetic region on the phenotype of interest. Within a genetic region, let 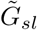 denote the quantile-transformed wavelet coefficient at scale *s* and location *l*. To test for association between the phenotype and the wavelet coefficient, we regress the wavelet coefficients on the phenotype Φ using a Normal prior. To adjust for covariates *C* that may be confounders in the GWAS, we incorporate the covariates into the regression models. The regression models for each scale and location are defined as follows:

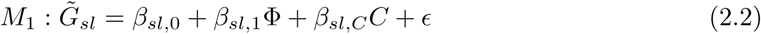

where *C* is a matrix of dimension *c* × 1 and *β*_*sl,C*_ is a matrix of dimension 1 × *c*, and *ϵ ∼ N* (0, 1). We compute the association parameters *β*_*sl*,1_ of the wavelet regression for each pair (*s, l*) using the closed form provided by Servin and Stephens (2007) for a normal prior with 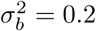. Under the null hypothesis, the 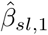 are normally distributed and have identical variance (due to the reverse regression). More specifically:

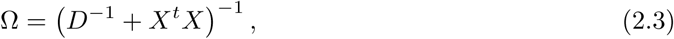

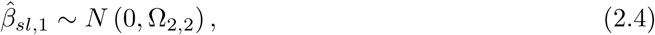

where *X* is the design matrix of dimension *K* × (2 + *c*). The design matrix includes the intercept, the phenotype, the confounding factors *c*, and the number of individual K. *D* is a diagonal matrix that includes the prior. Thus, 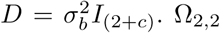 is the second element on the diagonal of Ω, given that we assume that the second column of *X* is the phenotype.

#### 2.3.1 Horizontal average evidence towards the alternative

For a given locus, a genetic signal might be assumed to occur in only a subset of the regression coefficients. Thus, the 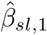 can be viewed as originating from a mixture of two Gaussian distributions, each representing a specific hypothesis:

- Under *H*_0_ the 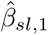 are distributed as 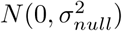.
- Under *H*_1_ some 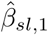 will be distributed as 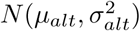.

To help identify a subset of *β* that convey the signal, we fit a mixture of the two Gaussian distributions to the collection of estimated 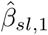, assuming mixture weights 1 − *π*_*alt*_ and *π*_*alt*_, respectively. Under the null hypothesis, the full mixture is not identifiable. In order to estimate 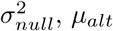, and 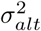 in all cases, and to ensure that the estimated 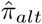 becomes small under *H*_0_, we constrain the Gaussian mixture fitting using a modified version of the EM algorithm with the restriction that 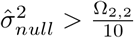 and 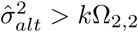, where *k* is of the order of 100.

After obtaining the estimates, we compute – for each beta coefficient – the posterior probability 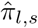 of *H*_1_ knowing *β*_*s,l*_ by

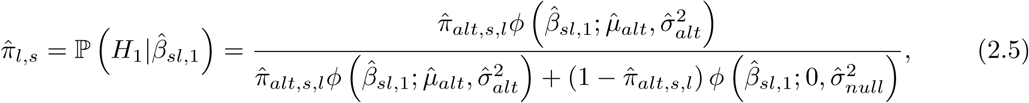

where *ϕ* (·; *µ, σ*^2^) is the density of a Gaussian distribution with mean *µ* and variance *σ*^2^. To reduce the influence of betas that most likely belong to *H*_0_, we propose a thresholding of the posterior probabilities 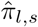 that decrease with sample size as well as with wavelet depth. Based on the work by Donoho and Johnstone (1994), we define the thresholded probabilities by

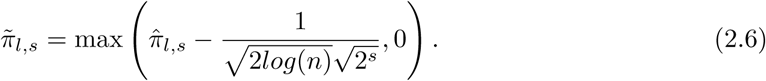

We later use the 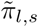 to localize the sub-regions of interest (for details, see Section 4.2 and Figure 5). Finally, we compute the horizontal average evidence towards the alternative by

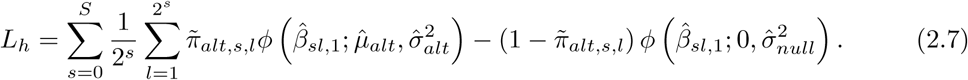

Note that, in contrast to Shim and Stephens (2015), our test statistic applies equal weight to each scale. In Shim and Stephens (2015), lower scales have more weight. As shown in the upper panel in Figure 2, the separation between the null and the alternative achieved by this test statistic (*L*_*h*_) alone is not optimal, with the resulting test power being low. See section 3 and the Supplementary Material for additional details.

#### 2.3.2 Inflation of the alternative hypothesis

To improve the separation of the null and alternative distributions, we extract two additional pieces of information from the 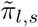. First, we compute the average proportion of associations per scale. The proposed test statistic is a weighted sum that applies the same weight to each scale. This can be seen as an alternative horizontal summary of the association:

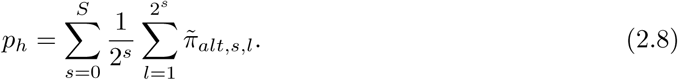

Second, we extract a vertical summary by considering sub-regions of the overall screened region. We divide the region of association into *S* − 1 sub-regions, where *S* is the maximum depth of analysis. We summarise the proportion of associations vertically, and for each average, we consider the positions that overlap with the sub-regions. For example, the first coefficient at scale 1 contributes half of the sub-region average of association.

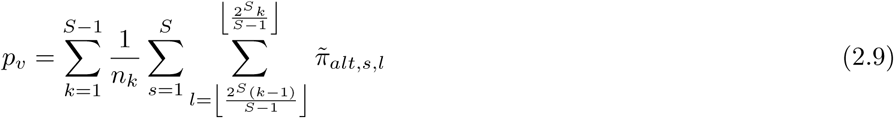

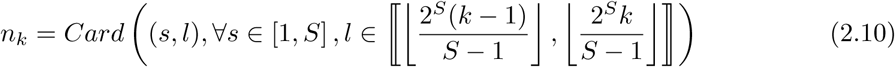

We use the new summaries of association to increase the separation between the null and the alternative by assuming that, under the alternative, *p*_*v*_ and *p*_*h*_ tend to be larger. We then build our full test statistic 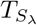, which requires calibration of the hyperparameter *λ*:

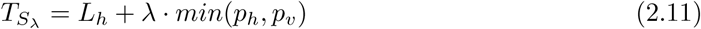

A larger *λ* would yield higher power if we assume that *p*_*v*_ and *p*_*h*_ tend to be larger under the alternative hypothesis. However, increasing *λ* can also change the shape of the null distribution. Assuming that the null distribution is normal, we use this as a fitting criterion to select the hyperparameter.

#### 2.3.3 Calibration of the hyperparameter and statistical significance

Our goal is to find the right balance between having as large of a *λ* value as possible while keeping the null distribution normal. As *min*(*p*_*h*_, *p*_*v*_) is not normally distributed (bounded distribution), the larger the *λ* is, the further 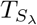 deviates from normality. In order to strike the right balance, we simulate *L*_*h*_ and *min*(*p*_*h*_, *p*_*v*_) under the null. Once simulated, we compute *L*_*h*_, *min*(*p*_*h*_, *p*_*v*_) for each simulation (10^5^ simulations in our case). Next, we fit a normal distribution on *L*_*h*_ and use this fit to generate the histogram of the p-values of the simulations for 1000 bins. We compute the number of counts in each bin and rank them by count (see Figure 3). We are particularly interested in the rank of the first bin, because an inflation of this bin would influence the false discovery rate. This procedure is repeated for increasing values of *λ*, and the search is stopped when a rank increase in the first bin is observed. We select the largest *λ* that results in the rank of the first bin to be equal to the rank of the first bin for *L*_*h*_, named *λ*\*. Finally, we use the normal fitting of 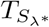 to perform the testing.

Given that the wavelet transform induces correlation between the 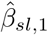, it is not possible to simulate them from a univariate normal distribution using their theoretical null distribution. One option is to compute the empirical covariance matrix of the 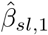 and simulate *β*_*sl*,1_ using this empirical null distribution. A second option is to simulate wavelets coefficients using random signals from a normal Gaussian distribution and then re-scale them to have a mean of zero and variance of one. Another possibility is to compute the covariance matrix of these wavelet coefficients and re-scale them using the theoretical null distribution of the 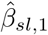. Similar results are achieved in both cases.

## 3. Simulations

### 3.1 Complex genetic signals

We performed simulations that correspond to complex genetic signals by combining real genetic data with a simulated phenotype. We used a previously identified locus for gestational age in the HARVEST dataset (see Magnus *and others* (2016)). Notably, we used the maternal genotype data from HARVEST spanning a region on chromosome 7 that covers base pairs 40504539 − 42504539 in the GRCH37-hg19 Human reference genome (Scherer, 2008). This region contains a total of 5209 SNPs in our data set. An example of the genetic variation in a single individual is displayed in Figure 1. We performed two sets of simulations to mirror so-called “local polygenic” effects. Each set of simulation considered different local polygenic effects by SNP location:

**Fig. 1.**
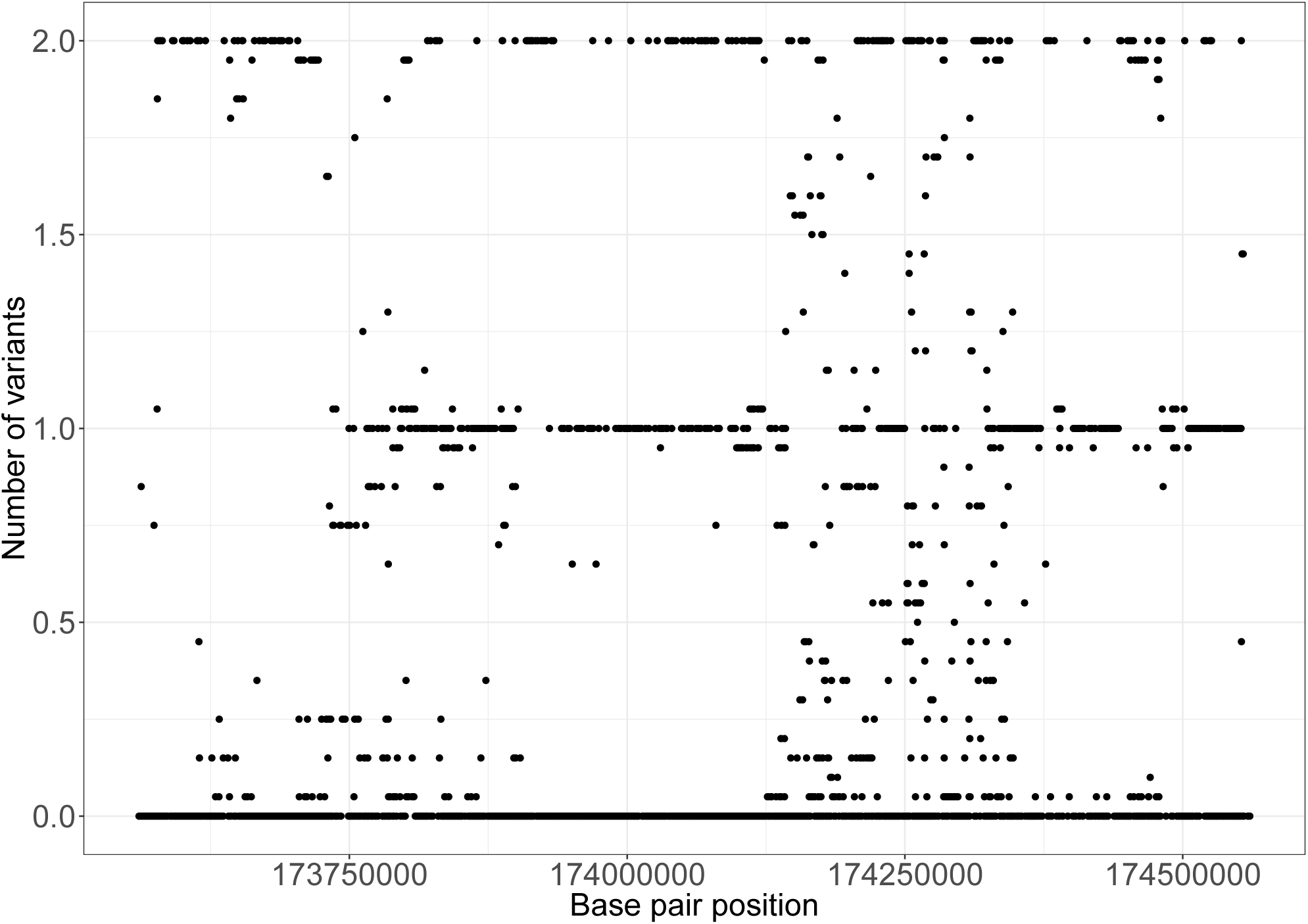
Genetic variation in one individual within a locus spanning two million base pairs (including 10000 imputed SNPs).

- *High LD Simulations*: we computed the correlation structure (LD) of the considered locus. We identified 28 small LD-blocks. In this simulation set-up, all the SNPs used in constructing the phenotype are selected within high LD regions. These simulations are engineered to mimic a diluted effect of SNPs within different LD blocks, also known as “block polygenic effect”, where each variant has a small additive effect.
- *Random LD Simulations*: all the SNPs (from 1 to 28) used in constructing the phenotype are taken uniformly at random from within the considered locus. These simulations are engineered to mimic a diluted effect of SNPs regardless of the LD structure, where each variant has a small additive effect.

In addition to the above simulations, we considered the following two models, each of which mimics different underlying locus effects.

- *Mono-directional model (MD)*: for each iteration, we randomly selected 1 to 28 SNPs. For each individual, we summed their SNP dosages within the selected set of SNPs to construct a score. On top of the individual scores, we added normally-distributed noise, scaled so that the genetic score explains 0.5% of the total phenotypic variance.
- *Random direction model (RD)*: the same setting as above, but the sign of the effect (positive/negative) for each SNP is random. In the mono-directional simulations, any additional variant may increase the level of the phenotype. This is not necessarily the case for random direction. Taken together, these simulations showcase the sensitivity of Wavelet Screening to the direction of the SNP coding.

The variance explained by a single SNP varies between 0.5%, which is typical for the top SNPs in a GWAS (Evan A. Boyle (2017)), to 0.018% – a level at which variants are not normally detected by the standard GWAS framework. We performed simulations for different sample sizes (1000, 5000, 10000). For each set-up, we performed 10000 simulations and ran Wavelet Screening on these using *c* and *d* coefficients. In addition, we performed 10000 simulations for no association, to assess the type 1 error for each sample size.

We performed 10^5^ simulations of *L*_*h*_ and *min*(*p*_*h*_, *p*_*v*_) for each simulation set-up. For each sample size, we searched for the hyperparameter (see 2.3.3). As displayed in Figure 2, there is a good match between the simulation and permutation distributions (we provide some examples in the Supplementary Material in Section 7 for other simulations and in Table 3 for type I error assessment).

**Fig. 2.**
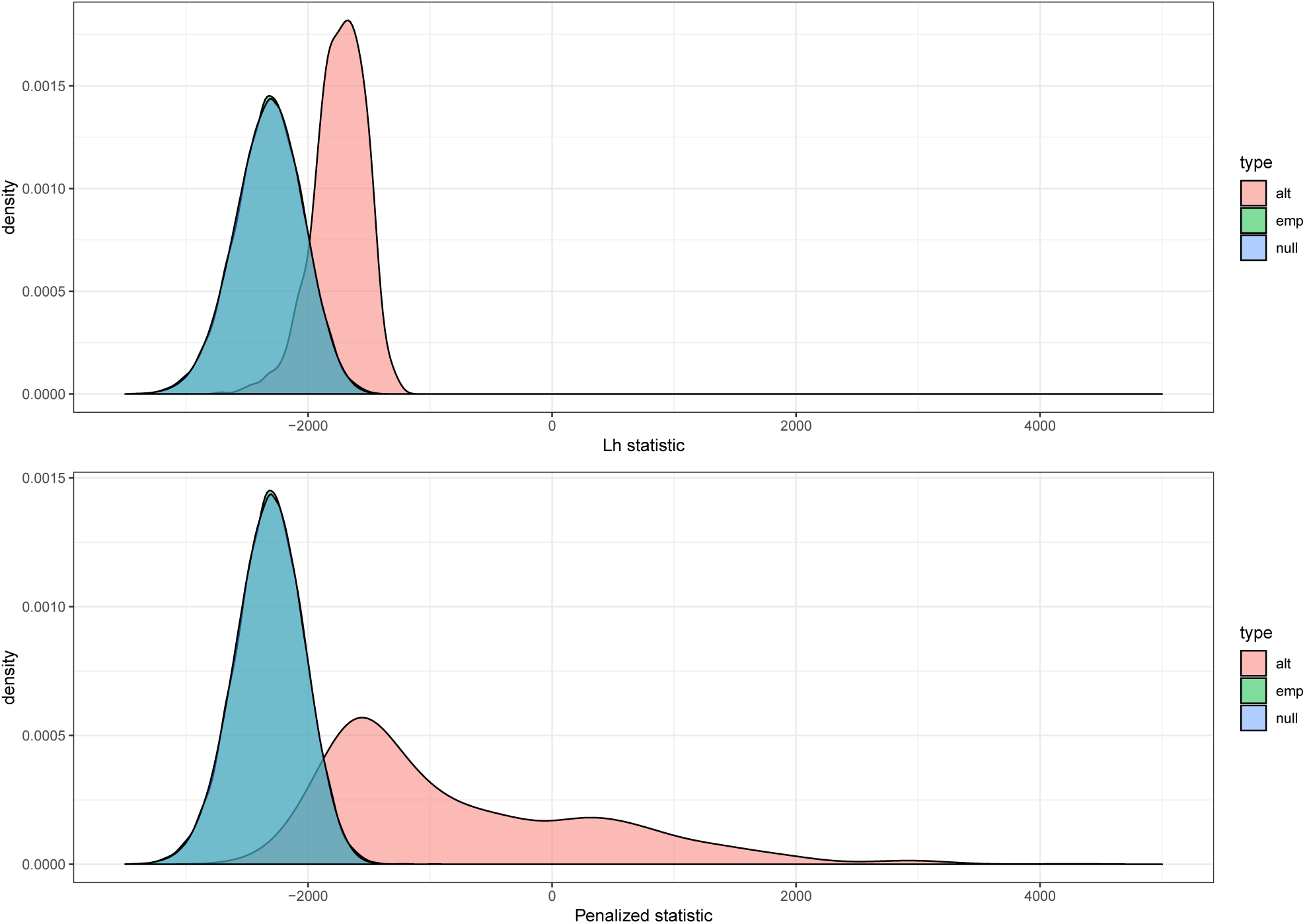
Horizontal average evidence towards the alternative. The simulated null distribution is shown in blue, the empirical null distribution in green, and the alternative distribution in pink.

**Table 1:** Power of the different methods depending on the number of components in the ‘Random LD Simulations’ (dilution effect). Numbers in bold show the maximum power for a given sample size and simulation set-up.

| Size | Model | Method | Significance | 1 | 2 | 3 | 4 | 5 | 6-10 | 11-15 | 16-20 | >20 |
| --- | --- | --- | --- | --- | --- | --- | --- | --- | --- | --- | --- | --- |
| 1000 | MD | WS | $1 \times 10^{-5}$ | **0.3** | **0.3** | **0.6** | **1.1** | **0.6** | **0.4** | **0.3** | **0.4** | **0.4** |
| 1000 | RD | WS | $1 \times 10^{-5}$ | **0.8** | 0.0 | 0.0 | 0.0 | 0.0 | **0.4** | **0.7** | 0.3 | 0.3 |
| 1000 | NA | SKAT-0 | $1 \times 10^{-7}$ | 0.0 | 0.0 | 0.0 | 0.0 | 0.0 | 0.0 | 0.0 | 0.0 | 0.0 |
| 1000 | NA | GWAS | $5 \times 10^{-8}$ | 0.0 | 0.0 | 0.0 | 0.0 | 0.0 | 0.0 | 0.0 | 0.0 | 0.0 |
| 5000 | MD | WS | $1 \times 10^{-5}$ | 9.4 | 10.6 | 8.2 | **6.0** | **4.0** | **3.6** | **2.4** | **2.3** | **2.5** |
| 5000 | RD | WS | $1 \times 10^{-5}$ | 9.4 | 8.5 | 7.1 | **5.1** | **4.6** | 2.2 | **1.4** | **1.0** | **1.7** |
| 5000 | NA | SKAT-0 | $1 \times 10^{-7}$ | 1.0 | 1.0 | 1.1 | 1.2 | 1.2 | 1.3 | 1.4 | 1.4 | 1.1 |
| 5000 | NA | GWAS | $5 \times 10^{-8}$ | **45.5** | **25.8** | **16** | 3.1 | 2.7 | 2.8 | 1.1 | 0.6 | 0.3 |
| 10000 | MD | WS | $1 \times 10^{-5}$ | 45.5 | 48.7 | 47.1 | 42.1 | **41.1** | **28.5** | **23.8** | **19.9** | **19.9** |
| 10000 | RD | WS | $1 \times 10^{-5}$ | 45.5 | 46.4 | 45.5 | 44.1 | 36.4 | 25.9 | **17.7** | **16.2** | **16.2** |
| 10000 | NA | SKAT-0 | $1 \times 10^{-7}$ | 8.3 | 7.2 | 8.6 | 8.6 | 8.4 | 10.4 | 10.3 | 9.8 | 7.9 |
| 10000 | NA | GWAS | $5 \times 10^{-8}$ | **100** | **81.8** | **75.0** | **68.8** | 30.6 | 28.1 | 15.5 | 9.6 | 3.3 |

**Table 2:** Power of the different methods depending on the number of components in the ‘High LD Simulations’ (dilution effect).

| Size | Model | Method | Significance | 1 | 2 | 3 | 4 | 5 | 6-10 | 11-15 | 16-20 | >20 |
| --- | --- | --- | --- | --- | --- | --- | --- | --- | --- | --- | --- | --- |
| 1000 | MD | WS | $1 \times 10^{-5}$ | **0.8** | **0.8** | **0.8** | **0.5** | **0.3** | **0.8** | **0.5** | **0.8** | **0.9** |
| 1000 | RD | WS | $1 \times 10^{-5}$ | **1.1** | **0.5** | 0.0 | **0.5** | **0.6** | **0.4** | **0.3** | **0.4** | **0.5** |
| 1000 | NA | GWAS | $5 \times 10^{-8}$ | 0.0 | 0.0 | 0.0 | 0.0 | 0.0 | 0.0 | 0.0 | 0.0 | 0.0 |
| 5000 | MD | WS | $1 \times 10^{-5}$ | 17.2 | 18.4 | **18.9** | **20.2** | **23.5** | **20.7** | **18.5** | **19.2** | **19.9** |
| 5000 | RD | WS | $1 \times 10^{-5}$ | 7.1 | 10.7 | 8.2 | **6.8** | **10.5** | **8.7** | **8.7** | **7.8** | **8.5** |
| 5000 | NA | GWAS | $5 \times 10^{-8}$ | 45.5 | 25.8 | 16 | 3.1 | 2.7 | 2.8 | 1.1 | 0.6 | 0.4 |
| 10000 | MD | WS | $1 \times 10^{-5}$ | 55.1 | 70.4 | **84.7** | **86.0** | **91.4** | **92.4** | **92.4** | **93.3** | **94.1** |
| 10000 | RD | WS | $1 \times 10^{-5}$ | 55.1 | 47.9 | 54.2 | 53.9 | **52.4** | **52.0** | **51.2** | **52.7** | **50.8** |
| 10000 | NA | GWAS | $5 \times 10^{-8}$ | 100 | 81.8 | 75.0 | 68.8 | 30.6 | 28.1 | 15.5 | 9.6 | 3.2 |

**Table 3:** Estimated Type I error for different sample sizes.

| $\alpha$ | 0.0500 | 0.0100 | 0.0010 | 0.0001 |
| --- | --- | --- | --- | --- |
| 1,000 | 0.0429 | 0.0065 | 0.0007 | 0.0002 |
| 5,000 | 0.0441 | 0.0068 | 0.0002 | 0.0001 |
| 10,000 | 0.0440 | 0.0075 | 0.0008 | 0.0006 |

We compared our method with the SNP-set (Sequence) Kernel Association Test (SKAT) (Ionita-Laza *and others*, 2013), which is a widely used method for regional-based association testing. SKAT aggregates individual SNP effects into a regional score to test for association. However, SKAT is not optimal for large regions. It is rather recommended for association testing in regions between 5 to 25 kb (Ionita-Laza *and others* (2013)). We thus performed our comparison by dividing our region of interest into sub-regions of 10 kb. We divided the genotype data into 5000 regions to perform a whole-genome screening using Wavelet Screening (see Section 4). Dividing these regions by 100 leads to a significance criterion of *p* = 10^−7^ for the SKAT method. In addition, as SKAT is more computer-intensive than our method, we used the power computation function of the R package *Power_Continuous_R* (Seunggeun Lee and Wu, 2017). This function uses simplified simulations to estimate the power of SKAT under different settings (see Lee *and others* (2011)). We used the argument r.corr = 2 to use SKAT-O, which is an optimised version for rare and common variants. In addition, we vary the proportion of causal SNPs according to the number of SNPs considered in the simulations for Wavelet Screening. Lastly, we make sure that for each set of simulations for a given proportion of causal SNPs, the total variance explained by all the loci is 0.5%. We use *BetaType* = *Fixed* and search for *b* (for *MaxBeta* = *b*) to obtain an explained variance of 0.5%. We mimic the Random Direction simulations by setting the argument *Negative*.*Percent* at 50%. For each sample size, type of effect, effect direction and number of SNPs, we performed 1000 simulations. Due to the heavy computational demands of SKAT, we only performed the *RandomLDSimulations*. This is because the user cannot specify the possible causal SNPs in *Power_Continuous_R*.

In Tables 1 and 2, *NA* means ‘not applicable’. As the standard single-SNP GWAS does not take sLD structure and local polygenicity into account, the only effect modelled here is the dilution effect. For SKAT-O, we performed simulations using the argument *Negative*.*Percent* (the proportion of negative effects) set at 50%. As we obtained similar results, we chose not to display them in Tables 1 and 2 but only specified the column *Model* with *NA* for SKAT-O.

### 3.2 Wavelet Screening improves discovery rate

The results in Tables 1 and 2 show that Wavelet Screening is a suitable alternative to GWAS when considering single-variant modelling. In both tables, we observed the dilution effect to be highly non-linear for the GWAS linear modeling, with a steep elbow-shaped curve. In contrast, the power for Wavelet Screening *d* and *c* coefficients decreases roughly linearly with the number of components in the score for *Random LD Simulation* and increases in case of *High LD Simulation*. SKAT-O has stable power under the dilution effect. However, it has lower power than the other methods and is a better alternative only when a very high number of variants have an effect on the phenotype. In addition, Wavelet Screening has higher power especially for smaller sample sizes. For *n* = 1000, none of the methods performed well. As seen in Tables 1 and 2, Wavelet Screening has non-zero power compared to SKAT-O and the GWAS based on single-variant modelling.

For *n* = 5000, Wavelet Screening outperformed all the other methods when at least four SNPs are considered in the simulation. For *n* = 10000, the GWAS based on single-variant modelling proved to be the best method when only 1-4 SNPs are considered. As shows in Table 2, Wavelet Screening has higher power in case of five SNPs or more within different LD blocks. For *Random LD Simulation*, Table 1 shows that the GWAS based on single-variant modelling is superior when up to five SNPs are considered and none of the SNPs are located within different LD blocks or are in LD with each other.

The power increase for a single SNP seen across Table 1 and 2 can be explained by the fact that the considered SNP for *High LD Simulation* is within an LD block. It is not necessarily the case with the *Random LD Simulation*. Wavelet Screening has higher power with the *c* coefficient. We conclude that Wavelet Screening improves the power of detecting an association substantially in case of *‘local polygenicity’*. As it is rare that only a single SNP is responsible for the association between a genotype and the phenotype, in a general setting, Wavelet Screening for the *c* coefficient would be more powerful than the standard GWAS based on single-variant modelling.

The type I error appears to be well-controlled by our procedure for the proposed simulation setting (see Table 3). As we only performed 10000 simulations under the null owing to computational burden issues, the results for *α* = 0.0001 are only indicative of the true error rate. For most situations, Wavelet Screening appears to be over-conservative, which might be due to our calibration criterion for *λ*\*.

## 4. Application

To test the utility of our new method, we performed a chromosome-wide association study of human gestational duration. Gestational duration is a complicated phenotype to study because of the combination of large measurement errors (*≈* 7 days, Nils-Halvdan Morken (2006)) and typically small genetic effects (*≈* 1.5 days (Zhang *and others*, 2017)). We used GWAS data on mothers from the Norwegian HARVEST study (Magnus *and others*, 2016) to replicate the lead SNPs reported in the largest GWAMA to date on gestational duration (Zhang *and others*, 2017). These lead SNPs are located on chromosome 5, near the gene for Early B cell factor 1 (*EBF1*). By using the same methodology as in Zhou and Guan (2017), as well as the same exclusion criteria for SNP and individuals, the lowest p-value obtained in our dataset was 2.8 × 10^−6^ for *n* = 8006, which is not statistically significant in the classic GWAS setting.

### 4.1 Definition of regions and choice of resolution

Although a typical GWAS can now interrogate millions of SNPs at a time, several chromosomal regions still have poor marker density, in particular near telomeres and centromeres, in regions of highly repetitive DNA, and in regions of low imputation quality. Most SNPs with low imputation quality are routinely discarded during quality control. Because we pre-process our data using an interpolation, we aim to avoid analysing purely interpolated regions. Our strategy entails including an additional criterion in the pre-processing step to exclude these types of regions. We propose studying regions of size 1 Mb (mega base-pairs), with a maximum distance of 10 kb between any two SNPs. Further, we define overlapping regions to avoid having a signal located at the very boundary of any one of the two regions. By applying these additional criteria, we excluded 18% of the SNPs and defined 248 regions on chromosome 5.

In addition to avoiding fully-interpolated regions, we also need to choose an appropriate depth of analysis for the wavelet decomposition. The precision of the wavelet coefficient depends on the number of non-interpolated points in a given region (Kovac and Silverman, 2000). As a rule of thumb, we propose to aim for 10 SNPs on average for each wavelet coefficient. Following this criterion, the median spacing between any given pair of SNPs was 202 base-pairs. This means that, if we divide each locus of 1 Mb into 2^9^ = 512 sub-regions, we would on average have 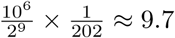 SNPs per sub-region.

### 4.2 Model and results

We applied Wavelet Screening to the gestational duration data set described above. In our modelling of gestational age, we included the first six principal components for each wavelet regression to control for residual population structure (Price *and others*, 2010). Next, we simulated *L*_*h*_ and *min*(*p*_*h*_, *p*_*v*_) 10^5^ times under the null using an empirical correlation matrix. Using these simulations and the steps described in Section 2.3.3, we obtained *λ*\* = 697999. We then fit a normal distribution on *L*_*h*_ + *λ*\**min*(*p*_*h*_, *p*_*v*_). This distribution was used to compute the p-values for each locus. These analyses identified two loci, but because we use half-overlapping windows, these two detected loci are the same. The discovered locus is depicted in Figure 5. Note that the associated p-value is smaller than the precision of the *qnorm* function in R version 3.5.2 (R Core Team, 2018) (i.e., below 10^−16^). The main SNP in the published GWAMA is located less than 1 Mb from the locus near *EBF1* that was detected by Wavelet Screening.

**Fig. 3.**
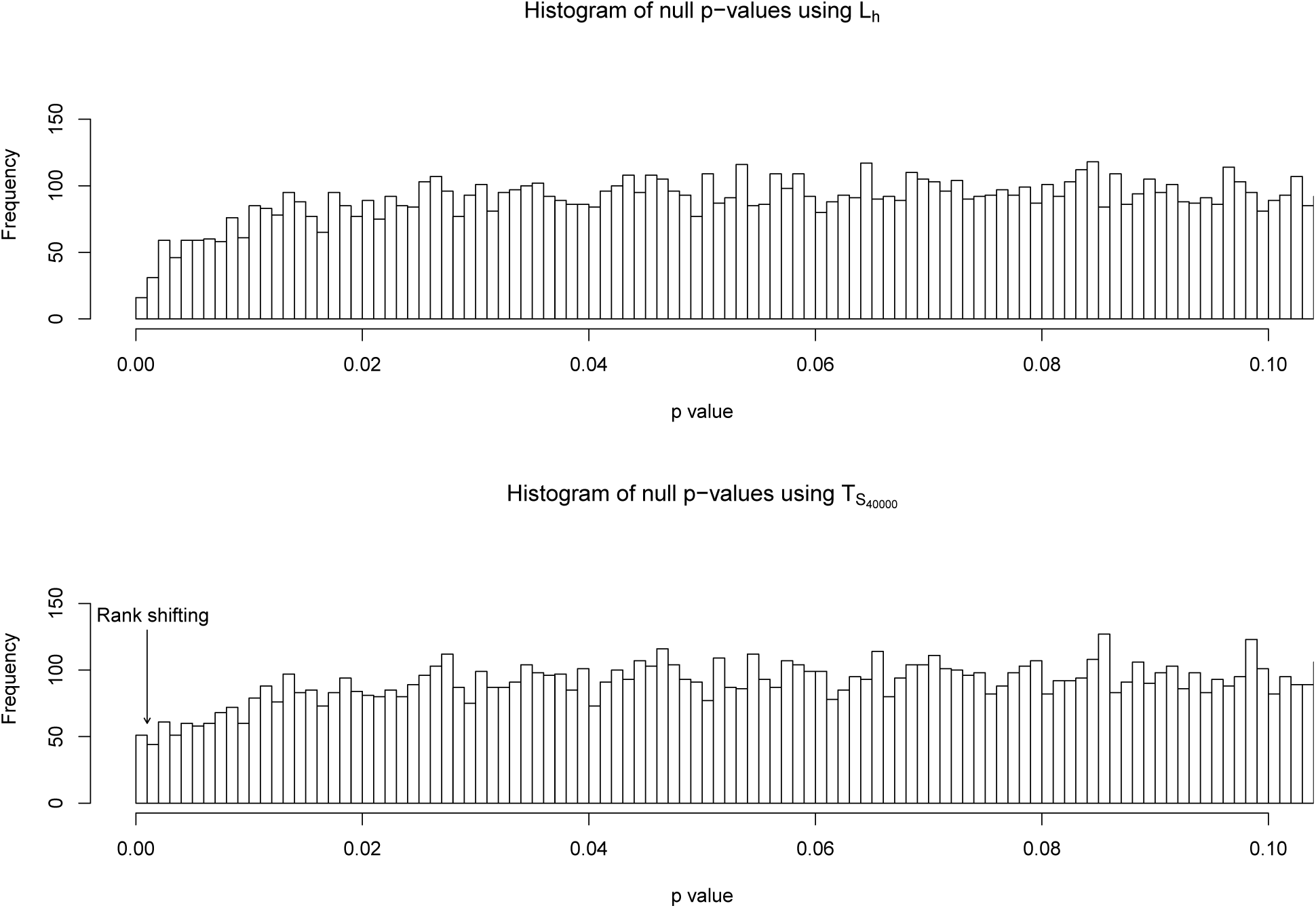
Rank shifting of the bin of interest.

**Fig. 4.**
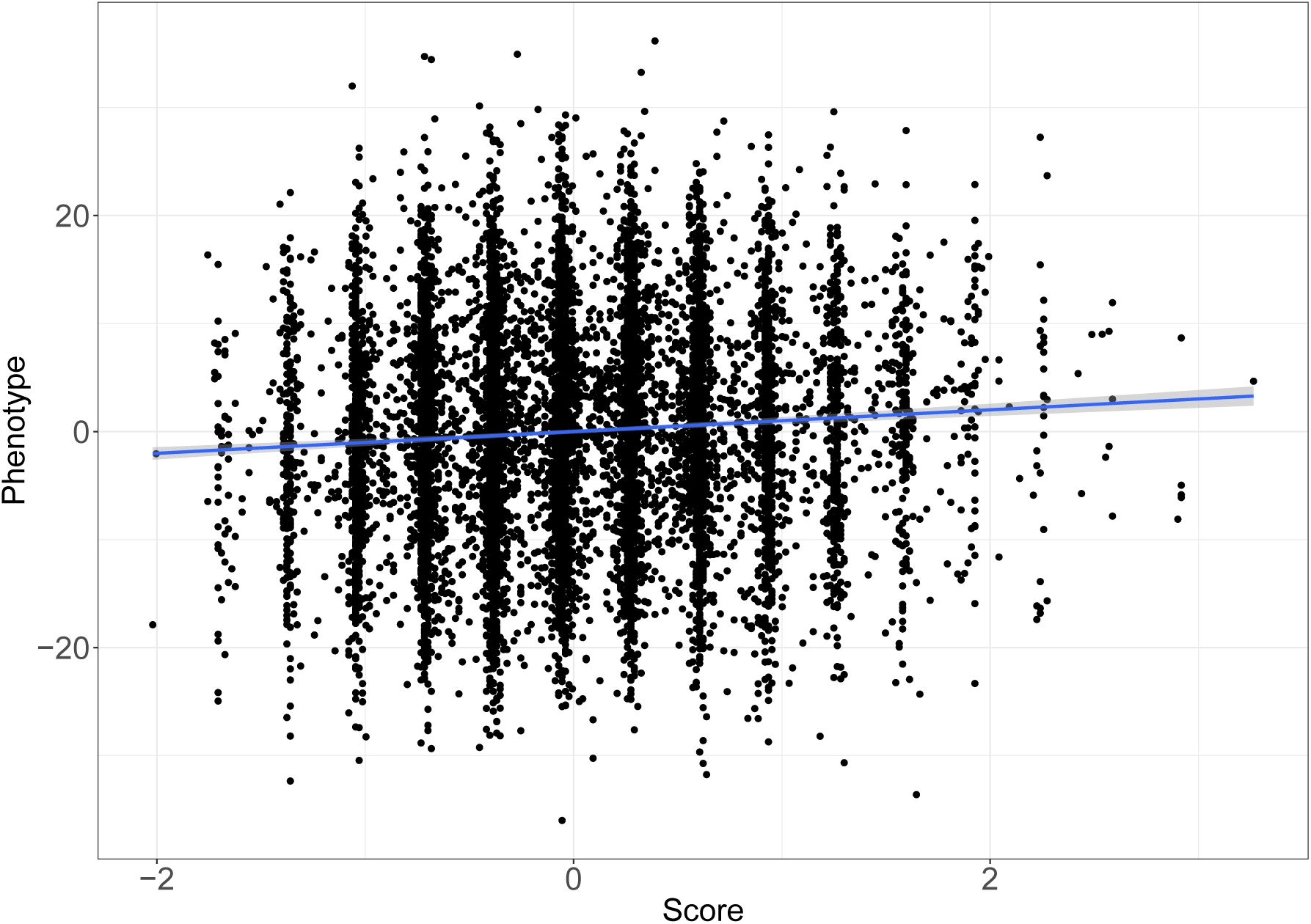
Simulated phenotype against the generated score (20 SNPs selected).

**Fig. 5.**
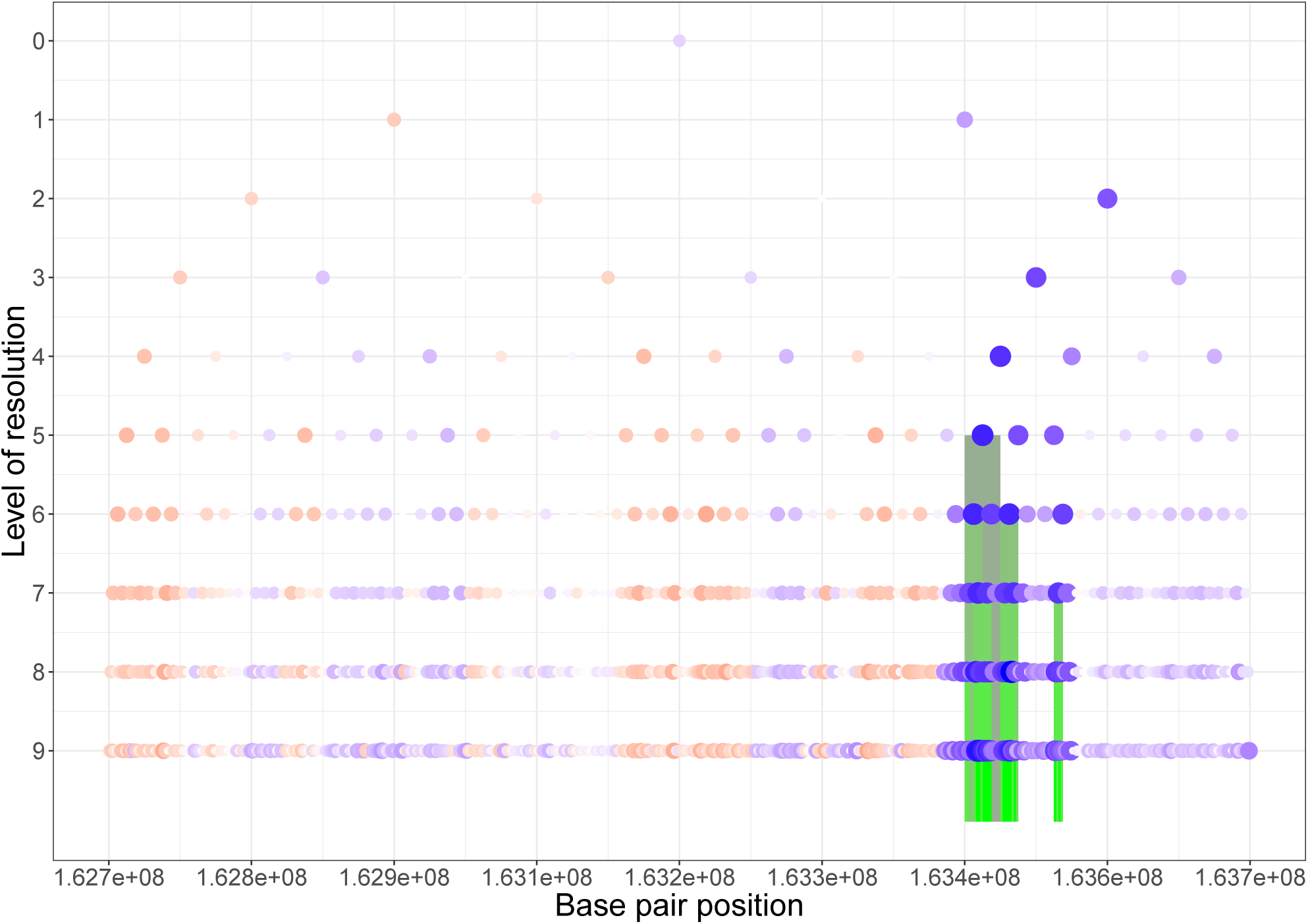
Locus discovered by Wavelet Screening. The dots of different sizes represent the absolute values of the estimated 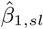; blue for negative, red for positive. The highlighted vertical bars represent 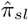 non-thresholded to zero.

We used the classic pyramidal wavelet decomposition representation to display the 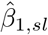 corresponding to each wavelet coefficient, with point size representing their absolute values and the colouring scheme representing their sign (blue for negative, red for positive). Furthermore, if a 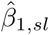 has an associated 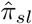 which is not thresholded to zero (see equation 2.5), we highlight the region corresponding to the wavelet coefficient using the colour-coding depicted in Figure 5.

## 5. Discussion

In this paper, we introduce Wavelet Screening as a novel and more powerful alternative to the classic GWAS methodology. It offers a more flexible modelling scheme than the standard single-point testing approach used in conventional GWAS and substantially improves the discovery rate. We acknowledge the empirical nature of this article, as most of the simulation set-ups indicate that the procedure is slightly over-conservative. This might be due to our calibration criterion for *λ*\*, or due to the shrinkage of the posterior probability. To alleviate some of these issues, we propose using a model under the null for 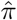.

Furthermore, we acknowledge the potential limitation of the coding scheme for assigning a SNP allele as either risk-conferring or protective. Minor alleles are conventionally coded as the risk alleles. When dealing with a large number of SNPs simultaneously, there is no definitive way of coding the alleles without prior knowledge of their true effects derived from the results of targeted functional analyses. When the risk allele is coded wrongly, the direction of the effect of the allele may be treated as random. In such a setting, Wavelet Screening would provide less power but would still be a better alternative to the single-SNP/variant modelling. Moreover, this coding limitation is ubiquitous and is present in all genotype-based regional tests that are not variance-based tests, such as SKAT-O or the Burden test (Ionita-Laza *and others*, 2013).

In future developments of Wavelet Screening, we plan to add functionalities to enable meta-analyses based on the use of summary statistics from different participating cohorts, akin to what is conventionally done using the METAL software (Cristen J. Willer and Abecasis, 2010). A meta-analysis should be straightforward to perform because the p-values across the different cohorts can be combined using Fisher’s method. However, it may be more appropriate to do a meta-analysis at the level of coefficients and then compute a new test statistic for the meta-analysis. In addition, we aim to adapt our method to include phenotypes on non-ordered scales, e.g., blood types or psychiatric phenotypes, which are usually analysed in a case-control fashion and not by multinomial regression because of computational and power issues. By exploiting reverse regression, we can include such phenotypes in the predictor matrix by coding them in a similar way to ANOVA. The modelling of the two hypotheses could be done using multivariate Gaussian, with one dimension per coefficient, instead of using simple univariate Gaussian. Further, by exploiting reverse regression, we can also easily adapt this method to the multiple-phenotype GWAS setting, also known as phenome-wide association studies or PheWAS (Zhonghua Liu, 2018).

Because of the complexity of the test statistics, it is difficult to infer directly how power would be influenced by the parameters (e.g., SNP distance, LD structure, the magnitude of the prior, etc.). Future work should focus on exploring power behavior under different scenarios, including sample size, percentage of variance explained, and unequally distributed effect between SNPs, among others. Lastly, our methodology is highly versatile in its applicability to various ‘omic’ data types. In future developments, we will investigate the feasibility of adding one more level of hierarchy to extend our method to multi-omic analyses.

## 6. Software

The Wavelet Screening method is distributed as an ***R*** package. In addition to the code, the package contains a data visualisation tool for exploring any association detected by Wavelet Screening. The ***R*** package is available at https://github.com/william-denault/WaveletScreening. To perform an analysis, the user only needs to specify one parameter (*σ*_*b*_). We also provide a detailed example using simulated data for how to use our package in the help function *wavelet_Screening*. Further, we show how to simulate *L*_*h*_ and *min*(*p*_*h*_, *p*_*v*_) under the null and the computation of *λ*\* from *L*_*h*_ and *min*(*p*_*h*_, *p*_*v*_), and, finally, the user can visualise the output of *Wavelet_Screening* as depicted in Figure 5 using the *plot_WS* function.

## Acknowledgements

This project was funded in part by the Research Council of Norway (RCN) (grant 249779). Additional funding was provided by the Bergen Medical Research Foundation (BMFS) (grant 807191) and the Research Council of Norway (RCN) through its Centres of Excellence funding scheme (grant 262700). The funders had no role in study design, data collection and analysis, decision to publish, or preparation of the manuscript. We are also grateful to Dr. Jonas Bacelis for his detailed comments on an earlier draft of the paper, and to Gatien Ricotier for his thorough review of the ***R*** package.

## Data access

The data used in the simulation section are publicly available. One can access these data by contacting the Norwegian Regional Ethical Committee (REK, *https://helseforskning.etikkom.no/*), and upon approval, data administrators at MoBa (*https://www.fhi.no/en/studies/moba/*) need to be contacted before secured access to the data can be provided.

## 7. Supportiong information

